# Investigating the roles of reflexes and central pattern generators in the control and modulation of human locomotion using a physiologically plausible neuromechanical model

**DOI:** 10.1101/2023.01.25.525432

**Authors:** Andrea Di Russo, Dimitar Stanev, Anushree Sabnis, Simon M. Danner, Jessica Ausborn, Stéphane Armand, Auke Ijspeert

## Abstract

Studying the neural components regulating movement in human locomotion is obstructed by the inability to perform invasive experimental recording. Neuromechanical simulations can provide insights by modeling the locomotor circuits. Past neuromechanical models proposed control of locomotion either driven by central pattern generators (CPGs) with simple sensory commands or by a purely reflex-based network regulated by state-machine mechanisms. However, the physiological interpretation of these state-machines remains unclear. Here, we present a physiologically plausible model to investigate spinal control and modulation of human locomotion. We propose a bio-inspired controller composed of two coupled central pattern generators (CPGs) that produce the rhythm and pattern and a reflex-based network simulating low-level reflex pathways and Renshaw cells. This reflex network is based on leaky-integration neurons, and the whole system does not rely on changing reflex gains according to the gait cycle state. The only component of the controller that maintains a state-machine mechanism is the balance controller of the trunk. The musculoskeletal model is composed of a skeletal structure and 9 muscles per leg generating movement in sagittal plane. After optimizing the open parameters, human locomotion replicating kinematics and muscle activation naturally emerged. Furthermore, when CPGs were not activated, no stable and physiologically plausible gaits could be achieved through optimization, suggesting the necessity of this component to generate rhythmic behavior without a state machine mechanism regulating reflex activation. The controller could reproduce a wide range of speeds from 0.3 to 1.9 m/s. The results also showed that the net influence of feedback on motoneurons during perturbed locomotion is predominantly inhibitory and that the CPG network provides the timing of motoneurons’ activation by exciting or inhibiting muscles in specific gait phases. The proposed bio-inspired controller could contribute to our understanding of locomotor circuits of the intact spinal cord and could be used to study neuromotor disorders.

## 1. Introduction

Limbs movements result from the complex interaction between brain centers, the spinal cord, and the musculoskeletal system [40]. The spinal network is essential in the control, coordination, and modulation of locomotion [27]. While there is direct evidence of a CPG in mammals and vertebrates [27, 1], the lack of direct experimental access in humans means that there is only indirect evidence [34]. Furthermore, sensory feedback pathways may play a major role compared to other mammals, and lower vertebrates in humans [29, 21, 22, 20]. Different studies suggested that the muscle activity observed during human locomotion may be controlled by five locomotor primitives that could be generated by rhythmic neural circuits [25, 10]. Hypothesizing these questions about the roles of different spinal components in controlling locomotion is challenging since we possess only partial knowledge of the interaction between the different subsystems involved in this process. On top of that, limited experimental access complicates the observation of the sub-components functions leading to difficulties in model validation. Computer simulations are necessary and have been proven useful in the past by evaluating the contribution of each control component by evaluating different models, and parameters [24, 15, 18, 36, 14, 3, 44].

Various neuromechanical models have been proposed in the past to address these questions. In 1995, Taga proposed a musculoskeletal system controlled by a neural rhythm generator composed of 7 pairs of neural oscillators and simple sensory-motor signals [45]. Successively, in 2001, Ogihara and Yamazaki developed a neural controller composed of motoneurons receiving inputs from a common CPG and reflexes from stretch and force receptors, where the spindle reflexes had inhibitory inputs to antagonist’s muscles [35]. In the context of locomotion controlled by CPG mechanisms, Aoi et al. (2010) constructed a CPG model based on a two-layered hierarchical network composed of a rhythm generator (RG) and a pattern formation (PF) layer. The RG model produced rhythmic information using phase oscillators and was regulated by phase resetting based on foot-contact gait events, whereas the PF model generated feedforward commands composed of five motor primitives based on the muscle synergies analysis performed by Ivanenko et al. [3, 25]. On the other hand, Geyer and Herr demonstrated that the kinematics and muscle activation observed in human locomotion could be reproduced without CPG commands by relying purely on sensory feedback activated at specific gait cycle phases [18]. A similar controller with partial modifications has been then proposed by Ong et al. [36]. In these studies, the activation of sensory responses in the gait cycle is regulated by a state-machine mechanism, hinting at the need for a more sophisticated circuit that controls the underlying reflexes. Other studies have integrated CPG commands on top of these purely sensory-based controllers, showing the benefits of rhythmic circuits in gait modulation [15, 48].

In this study, we propose a novel bio-inspired controller composed of a feedforward network inspired by Aoi et al. consisting of two CPGs that produce the locomotor rhythm and patterns and a new physiologically plausible implementation of spinal reflexes based on neurophysiological studies in locomotion [51, 49] without relying on any state machine mechanism. This network controls 9 muscle actuators generating torques in a previously assessed musculoskeletal model [13]. The performance of this controller in replicating the behavior of human locomotion and its modulation are investigated and compared with previous experimental and neuromechanical studies. In addition, we investigate the performance of the sensory feedback controller alone to verify whether it is possible to generate human walking behavior with a purely reflex-based controller without relying on state machine mechanisms and to verify the benefit of CPG mechanisms. Finally, we examine the contribution given by pattern generation and reflex circuits to the motoneurons at slow, intermediate, and fast speeds performing a correlation analysis to identify possible parameters responsible for speed modulation. With these experiments, we aim to address the following questions:

- What is the role of CPGs and reflex circuits in the generation of muscle activation in human locomotion?
- Can low-level feedback circuits produce stable locomotion without a CPG or statemachine?”
- Is the contribution of these two neural components changing with increasing of gait speed?

Our results show that the reflex rules implemented in previous models [18, 36] could be reproduced into less abstract models of neural circuits. The insights given by the proposed controller suggest that spinal reflexes alone could not reproduce rhythmic locomotion without a state machine mechanism regulating the activation of reflexes in specific phases of the gait cycle. CPG networks appear to play the role of state machines in previous models and to be necessary to promote muscle activation in specific gait cycle phases. In addition, the modulation of CPG frequency seems necessary to modulate step duration. The modulation of either reflexes, CPG network, or both could generate gaits in a wide speed range, highlighting the high level of versability of the neurospinal control of human locomotion.

## 2. Methods

This study used the SCONE software simulation framework [17], which was extended to implement and optimize the new spinal model generating gait simulations of 10 s. The SCONE simulation comprises the following four blocks:

- An OpenSim musculoskeletal model.
- A controller.
- A cost function composed of several locomotion metrics
- An optimizer that optimizes the initial conditions and controller parameters to minimize the cost function.

### 2.1. Musculoskeletal model

The musculoskeletal model (Figure 5) has the skeletal structure presented by Delp et al. [13] with a height of 1.8 m and weight of 75.16 kg. The model is constrained in the sagittal plane and has a total of nine degrees of freedom (DoFs): a 3-DoFs planar joint between the pelvis and the ground and 3 rotational DoFs per leg: hip flexion/extension, knee flexion/extension, and ankle dorsiflexion/plantarflexion. Three spheres per leg are used as contact models with the ground: one of radius 5 cm at the calcaneus and two of radius 2.5 cm at the toes. The musculoskeletal model is actuated by nine Hill-type muscle-tendon units per leg: gluteus maximus (GMAX), biarticular hamstrings (HAMS), biceps femoris short head (BFSH), rectus femoris (RF), iliopsoas (ILPSO), vasti muscle group (VAS), gastrocnemius (GAS), soleus (SOL), and tibialis anterior (TA).

### 2.2. Controller

Muscle activation is regulated by the excitation provided by the motoneurons. The motoneurons are stimulated or inhibited by the different components of the bio-inspired controller: the balance controller of the trunk and the spinal network, composed of the CPGs and spinal reflexes. The balance controller and the CPGs are modeled at an abstract level. Indeed, the former provides balance inputs in specific phases of the gait cycle with Proportional Derivative (PD) controllers, and the latter is composed of two abstract oscillators generating primitive patterns. By contrast, the spinal reflexes are modeled at a lower level of abstraction and are structured in different leaky integrator neurons divided into three types: somatosensory neurons (*S Ns*), interneurons (*INs*), and motoneurons (*MNs*). The overall structure of the controller is reported in Figure 2. The balance controller of the trunk regulates only the activation of hip muscles in specific phases of the gait cycle, whereas CPGs and spinal reflex controllers provide inputs to all muscles and are not regulated by any state-machine mechanism. We chose to maintain the state machine for the balance controller in order to simplify the balance control since our main goal is the simulation of locomotor movement. A physiological neuromechanical model of trunk balance control is a complex task that is outside the scope of this study. Muscle excitation is triggered by the motoneuron output with values between 0 and 1 since motoneurons can only provide excitation to muscle fibers and cannot have negative outputs. To keep a reasonable level of abstraction and complexity, we will assume that the neuron’s output *n*_*o*_*utput* follows the dynamics of a leaky integrator:

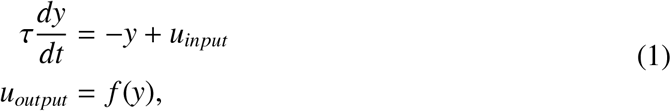

where *y* is the neuronal response, *u*_*input*_ is the neural input, τ the time constant (typically 0.01), *u*_*output*_ the output of the neuron, and *f* the activation function. The activation function used for motoneurons is the min-max operator (*f* (*x*) = *min*(*max*(0, *x*), 1)), and the neural input is defined as:

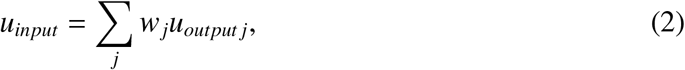

where *w*_*j*_ is the weight associated with the *j*^*th*^ connection, and *u*_*output j*_ the output of the *j*^*th*^ neuron. The motoneuron receives inputs from the CPGs’ network (*u*_*CPGs*_) and the reflex circuit (*u*_*re f lexes*_). These inputs are integrated according to equations 1 and 2 and generate the motoneuron output *m*_*output*_. ILPSO, GMAX, and HAMS also receive inputs from the balance controller (*u*_*balance*_). To avoid the activity of the balance controller from being inhibited by the other neural circuits possibly preventing the correct balance of the trunk, *u*_*balance*_ is not integrated into the motoneuron dynamics, and the final motoneuron output for hip muscles 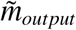 is defined by the following equation:

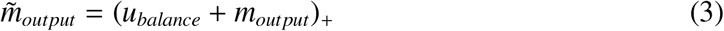

where *m*_*output*_ is the motoneuron output resulting from the integration of *u*_*CPGs*_ and *u*_*re f lexes*_ and *u*_*balance*_ represents the balance controller effect on the hip muscles. The operator ()_+_ represents only the positive part of the selected signal. The amplitude of all components is regulated by the controller’s parameters tuned by the optimization algorithm. The muscle activation *a* responds to the excitation *m*_*output*_ (or 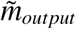 _*output*_ for hip muscles) as defined by Thelen et al. [46]. The following sections will describe in detail how each neural input is computed (*u*_*balance*_,*u*_*CPGs*_, and *u*_*re f lexes*_).

**Figure 1:**
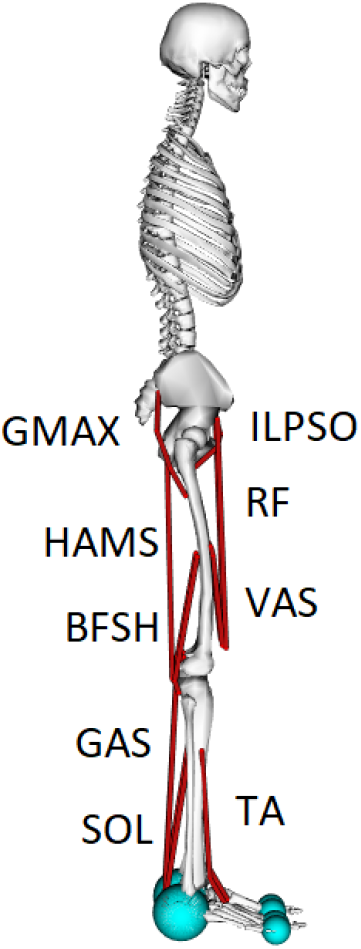
Musculoskeletal model used to study human locomotion. The model is constrained in the sagittal plane and has 9 DoFs: hip and knee flexion/extension, ankle plantar/dorsal flexion for each leg, and a 3-DoFs planar joint between the pelvis and the ground. Movements are generated by the activation of 9 muscles per leg: gluteus maximus (GMAX), biarticular hamstrings (HAMS), biceps femoris short head (BFSH), rectus femoris (RF), iliopsoas (ILPSO), vasti (VAS), gastrocnemius medialis (GAS), soleus (SOL), and tibialis anterior (TA).

**Figure 2:**
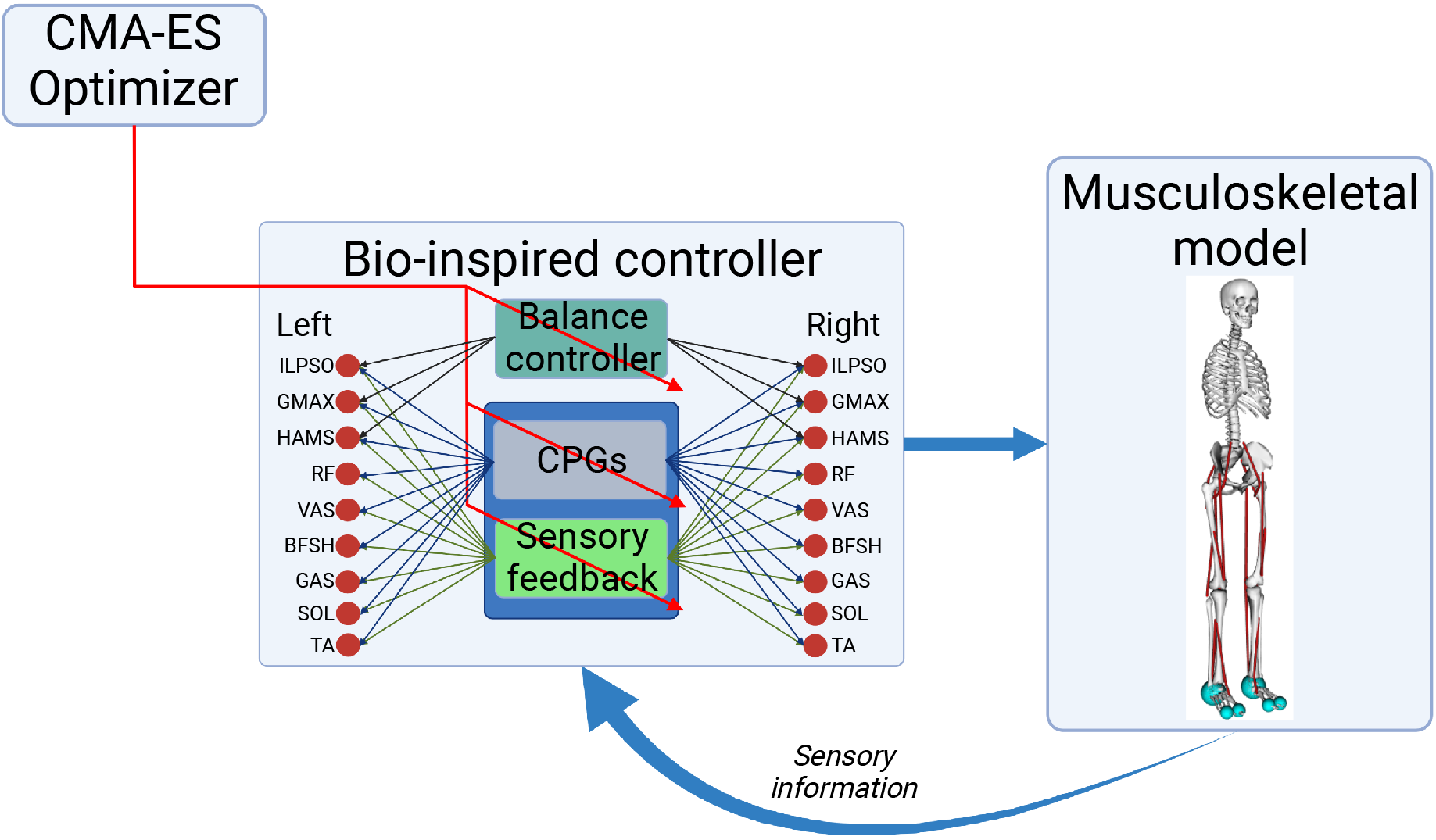
Control diagram: the bio-inspired controller is composed of the balance controller of the trunk and the spinal controller divided into CPGs and spinal reflexes. The balance controller aims at keeping the balance of the trunk by stimulating the hip muscles’ motoneurons, whereas CPGs and spinal reflexes generate rhythmic behavior stimulating all muscles’ motoneurons. Reflexes and CPGs are integrated by motoneurons, whereas balance inputs are summed separately. (Created with BioRender.com)

#### 2.2.1. Balance controller of the trunk

The balance controller is the one proposed by Ong et al. [36], and it is the only controller part where a state-machine mechanism is present. A proportional derivative control strategy (PD) is used to activate the hip muscles balancing the forward lean angle of the trunk. ILPSO, GMAX, and HAMS receive inputs from the balance controller during the stance phase. The excitation given by the balance controller to the hip motoneurons is described in equation 4.

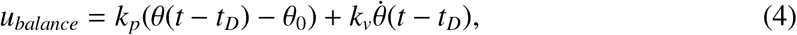

where *k*_*p*_ and *k*_*v*_ are the proportional and derivative controller’s gains, and the constant θ_0_ is the desired forward lean angle regulating the proportional feedback of the actual forward lean angle θ. *t*_*D*_ represents the time delay, corresponding to *t*_*D*_ = 5*ms* for the hip muscles. The balance controller has a total of 9 parameters.

#### 2.2.2. Central Pattern Generators (CPGs)

The central pattern generators (CPGs) were implemented as two coupled oscillators (one per side) composed of rhythm generator, and pattern formation layers [41, 31] inspired by the work of Aoi et al. [3, 2]. The rhythm generator dictates a period command synchronized with the environment through afferents triggered by the heel-strike event. Based on Aoi’s model, two coupled differential equations govern the CPG dynamics:

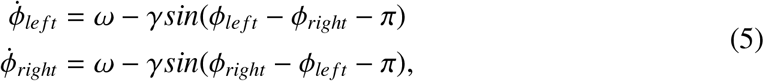

where ϕ_*le f t*/*right*_ denotes the phase of each leg, ω(*t*) is the basic angular frequency, and γ is the coupling constant.

The differential equation contains events that reset ϕ_*le f t*/*right*_ when the leg touches the ground in order to synchronize the CPGs’ phases with the environment. This is the only feedback mechanism present in the CPGs model, and it is described in equation 6:

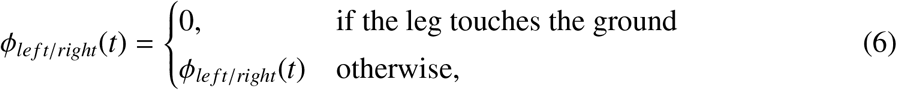

In our simulations, the angular frequency ω has a constant value and represents one of the parameters under optimization.

The pattern formation layer is composed of phase-dependent primitive patterns. Each pattern resembles a bell-shaped waveform with a defined width that can be centered around a specific phase value and is implemented as a raised-cosine function:

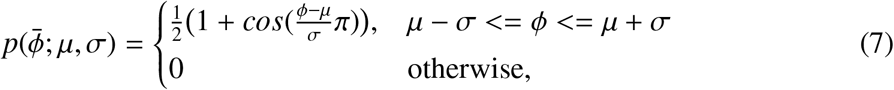

where 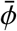 is the normalized gait phase, µ is the value corresponding to the peak of the bell shape, and σ is the half-width of the curve. The pattern formation layer is composed of five primitives of the same half-width and centered at different times of the gait phase (Figure 3a):

**Figure 3:**
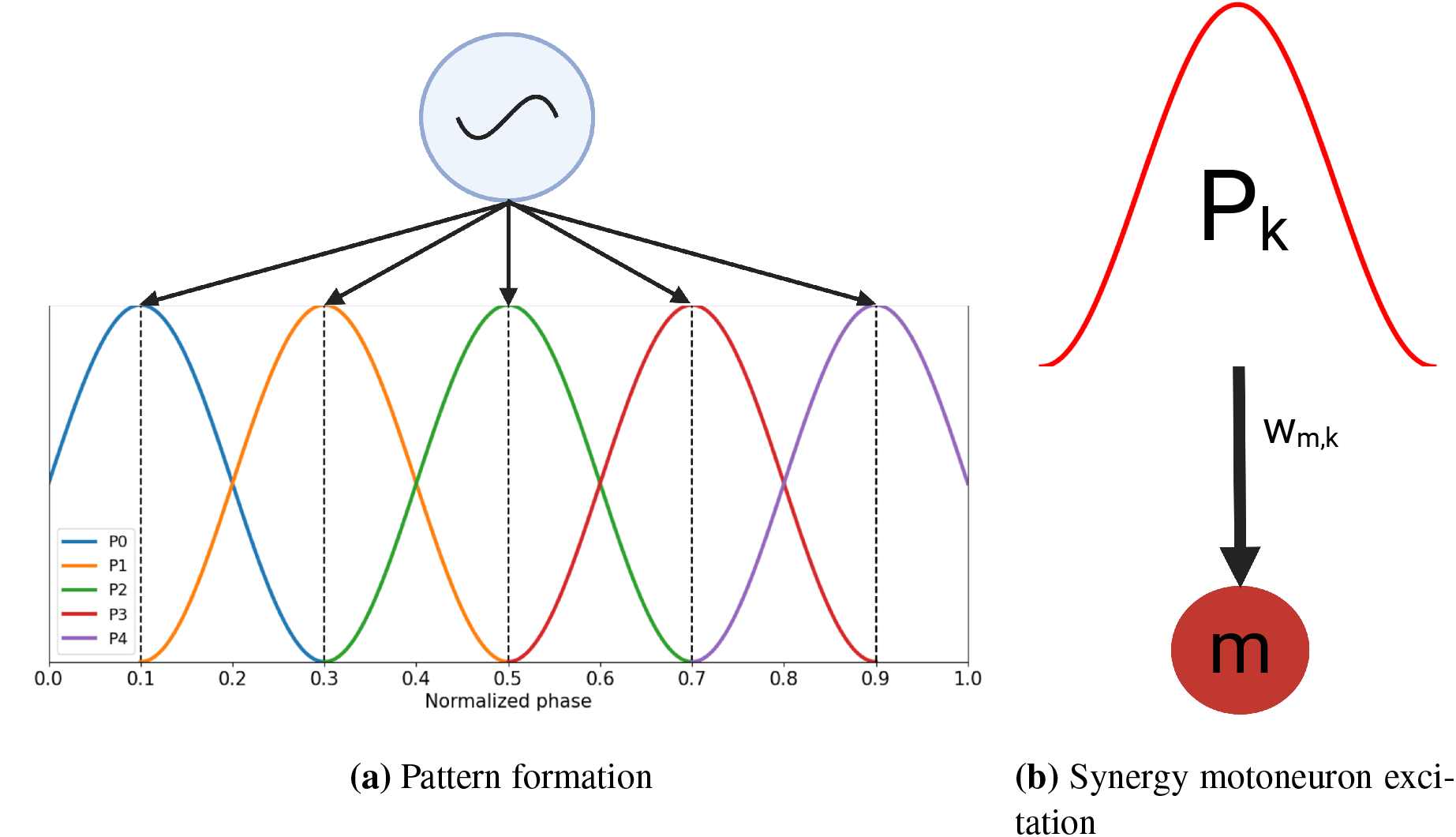
CPG structure: 3a) The CPG generates five bell-shaped primitives centered at different times of the gait cycle. 3b) Each *k*-pattern stimulates all the *m*-motoneurons depending on the assigned weight *w*_*m,k*_ that can be positive or negative. (Created with BioRender.com)

- P0: *µ* = 0.1, *σ* = 0.2
- P1: *µ* = 0.3, *σ* = 0.2
- P2: *µ* = 0.5, *σ* = 0.2
- P3: *µ* = 0.7, *σ* = 0.2
- P4: *µ* = 0.9, *σ* = 0.2

The choice of modeling the CPG network as the generation of five locomotor primitives derives from the observations done in past experimental studies where five bell-shaped synergies active at different phases of the gait cycle were identified in human studies [25, 26]. Each motoneuron receives a weighted neural excitation or inhibition *u*_*CPGs*_ from all primitive patterns (Figure 3b) according to the following equation:

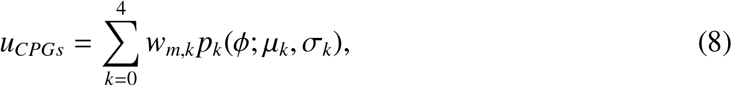

where *w*_*m,k*_ is the weight parameter of the pattern *k* to the motoneuron *m* to be determined through optimization. The total number of parameters to optimize corresponds to 5 weights per pattern to each specific muscle and the oscillatory frequency. Therefore, the number of parameters for the CPG network is 48. The possible values assigned to *w*_*m,k*_ are [-1:1].

#### 2.2.3. Spinal reflexes

To implement a physiologically realistic model of sensory-motor control in human locomotion, we model and investigate five spinal reflexes:

- Ia afferents provide monosynaptic excitation to motoneurons innervating the same muscle and disynaptic inhibition mediated by Ia inhibitory interneurons to antagonistic motoneurons (Figure 4a) and model the velocity-dependent response to stretch [9].
- II afferents provide disynaptic excitation to motoneurons innervating the same muscle and disynaptic inhibition to antagonistic motoneurons mediated by excitatory and inhibitory interneurons, repectively(Figure 4b). This reflex models the excitatory role of group II afferents [30] responding to changes in muscle length during stretch.
- Ib afferents provide disynaptic inhibition to motoneurons innervating the same muscle mediated by inhibitory interneurons.These interneurons reciprocally inhibits with antagonistic Ib-interneurons (Figure 4c). This reflex is triggered by the Golgi tendon organs and it is introduced to protect muscles when large forces are detected [7]. Additionally, Ib afferents provide dysinaptic excitation to extensor motoneurons innervating the same muscle mediated by excitatory interneurons. These connections model the positive force feedback reversal commonly observed in experimental studies [39, 19, 41] (Figure 4d).
- Renshaw cells are inhibitory interneurons providing inhibitions to motorneurons and Iainterneurons innervating the same muscle. Additionally, these cells reciprocally inhibits with antagonistic Renshaw cells [50] (Figure 4e). Renshaw cells are activated by motoneurons innervating the same muscle though synaptic excitation inhibiting these motoneurons when a large activity is detected.

The spinal sensory feedback network is composed of three types of leaky integrator neurons: somatosensory neurons (*S Ns*), interneurons (*INs*), and motoneurons (*MNs*). Each of these neurons model the activities of neural populations in the physiological spinal cord. *INs* have the same properties of *MNs* responding to the dynamics described in equations 1 and 2 and with the same activation function *f* (*x*). *S Ns* instead presents a rectifier function (*f* (*x*) = *max*(0, *x*)) as activation function, and also the neural input is slightly different:

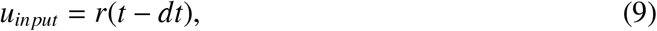

where *r*(*t* − *dt*) is the receptor function and *dt* the delayed value of the receptor. Transmission delays are known and can be determined according to the proximity of the receptors [15]. The expressions of the receptors follow the equations:

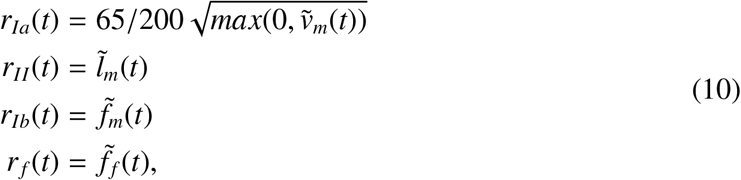

Where 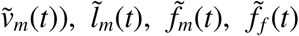, are respectively the normalized quantities for contraction velocity, muscle length, muscle force, and cutaneous forces due to ground-foot contact. We choose to consider the normalized quantities to be able to easily scale for different muscles with different values of length and strength. Here, the expression for *r*_*Ia*_ was inspired by Prochazka [38] and modified such that only lengthening 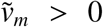 triggers a response while ignoring length and activity-dependent terms. We deliberately simplified these expressions because we wanted to capture the general trend and prevent an excessive number of physiological parameters. In Figure 4, we present the primitive reflex pathways that govern the connectivity within a single spinal cord segment. These rules are used to build the topological network by assuming that muscles can be categorized as agonists (A), antagonists (N), and extensors (E) or flexors (F).

**Figure 4:**
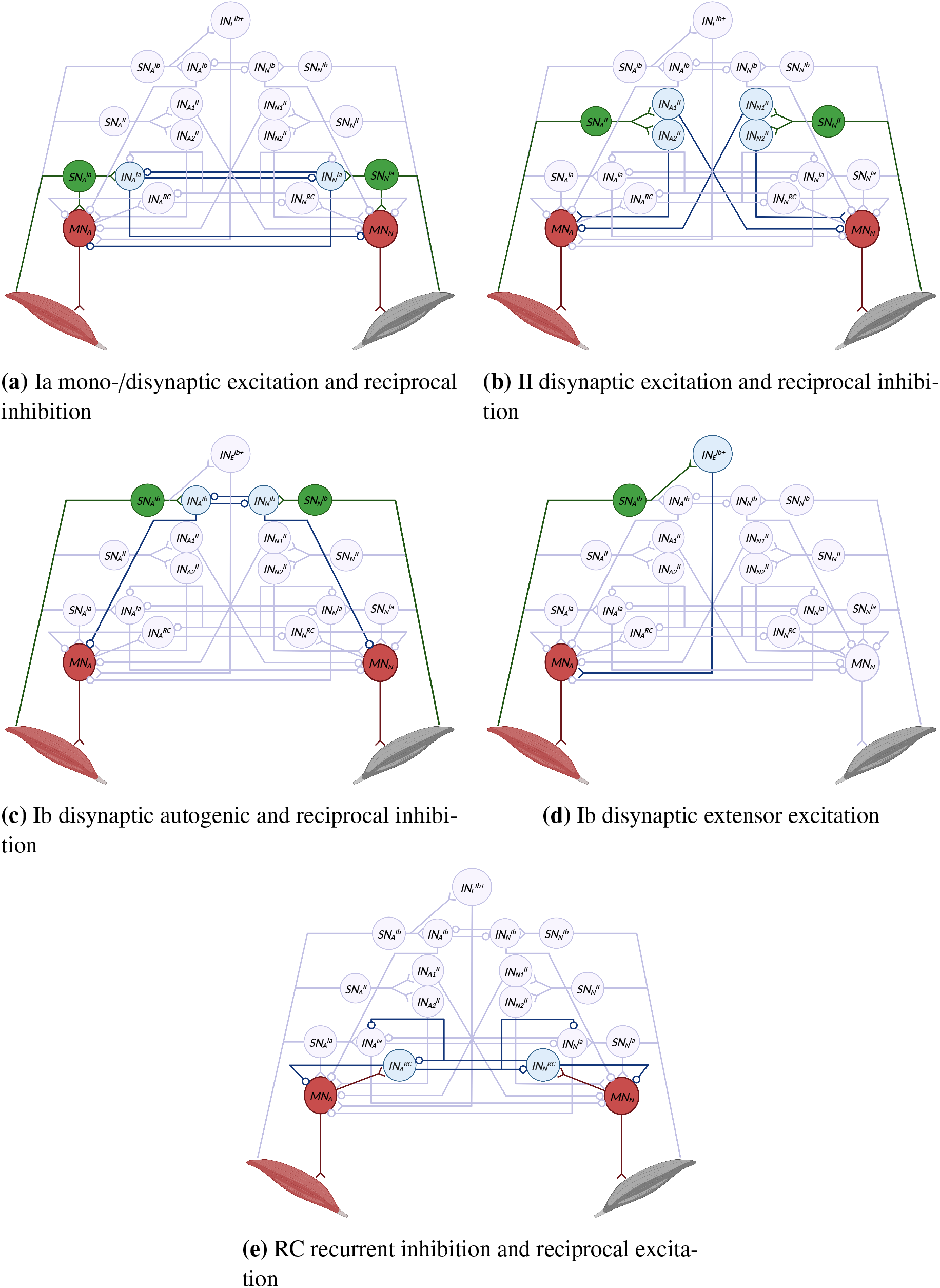
Reflex pathways. Green, blue and red stand for somatosensory (*S N*_*A*/*N*_), inter (*IN*_*A*/*N*_), and motor neurons (*MN*_*A*/*N*_), respectively. The connection tip *o* stands for inhibition while < is for excitation. Subscript letters A, N, and E denote agonist, antagonist, and extensor muscles, respectively. The rules are repeated for all antagonist muscles. (Created with BioRender.com)

The relation between agonist and antagonist muscles defines the mutual inhibitions described in Figure 4a, 4c, 4b, and 4e. In addition, a muscle can be defined as extensor of flexor. In case it is an extensor muscle, the additional connections of Ib disynaptic extensor facilitation described in Figure 4d are included. Some bi-articular muscles can be considered both extensors and flexors since they have different effects on different joints and the Ib disynaptic extensor facilitation is included also in this case. Table 1 describes the relations among agonist and antagonist muscles assigned in our models. Accounting for all the weighted connections, the sensory feedback controller has a total of 183 parameters.

**Table 1:**
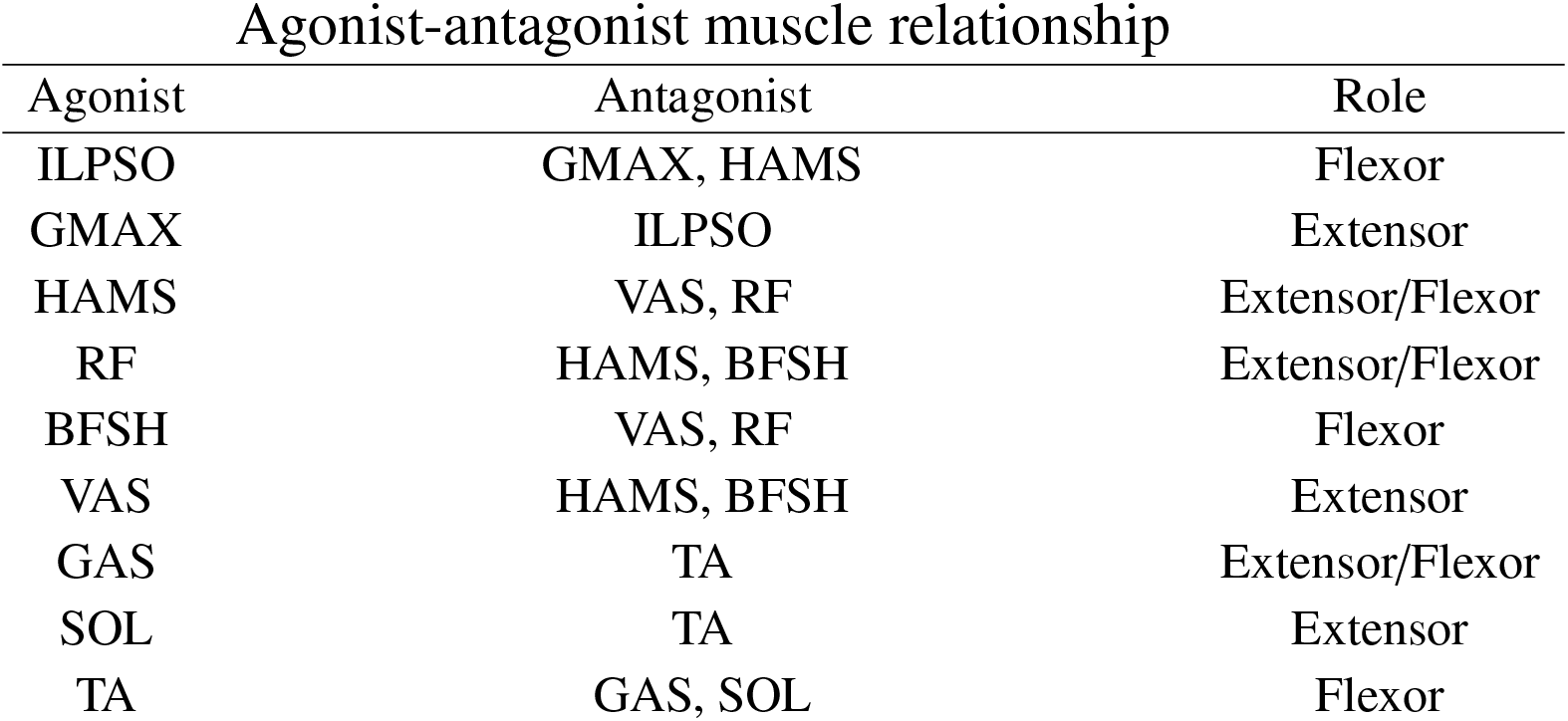
Agonist-antagonist relationship among muscles modeled. Each antagonist relationship implies the corresponding reciprocal inhibition of Ia, II, and Ib connections, and the reciprocal excitation connections of RC. The table specifies whether the agonist is considered an extensor, which includes the disynaptic excitation from Ib+, or flexor.

| Agonist-antagonist muscle relationship |  |  |
| --- | --- | --- |
| Agonist | Antagonist | Role |
| ILPSO | GMAX, HAMS | Flexor |
| GMAX | ILPSO | Extensor |
| HAMS | VAS, RF | Extensor/Flexor |
| RF | HAMS, BFSH | Extensor/Flexor |
| BFSH | VAS, RF | Flexor |
| VAS | HAMS, BFSH | Extensor |
| GAS | TA | Extensor/Flexor |
| SOL | TA | Extensor |
| TA | GAS, SOL | Flexor |

Finally, Figure 5 shows the whole spinal network implemented between agonist and antagonist muscles including the reflex pathways and CPGs inputs. On top of this network, ILPSO, GMAX, and HAMS also receive inputs from the balance controller.

**Figure 5:**
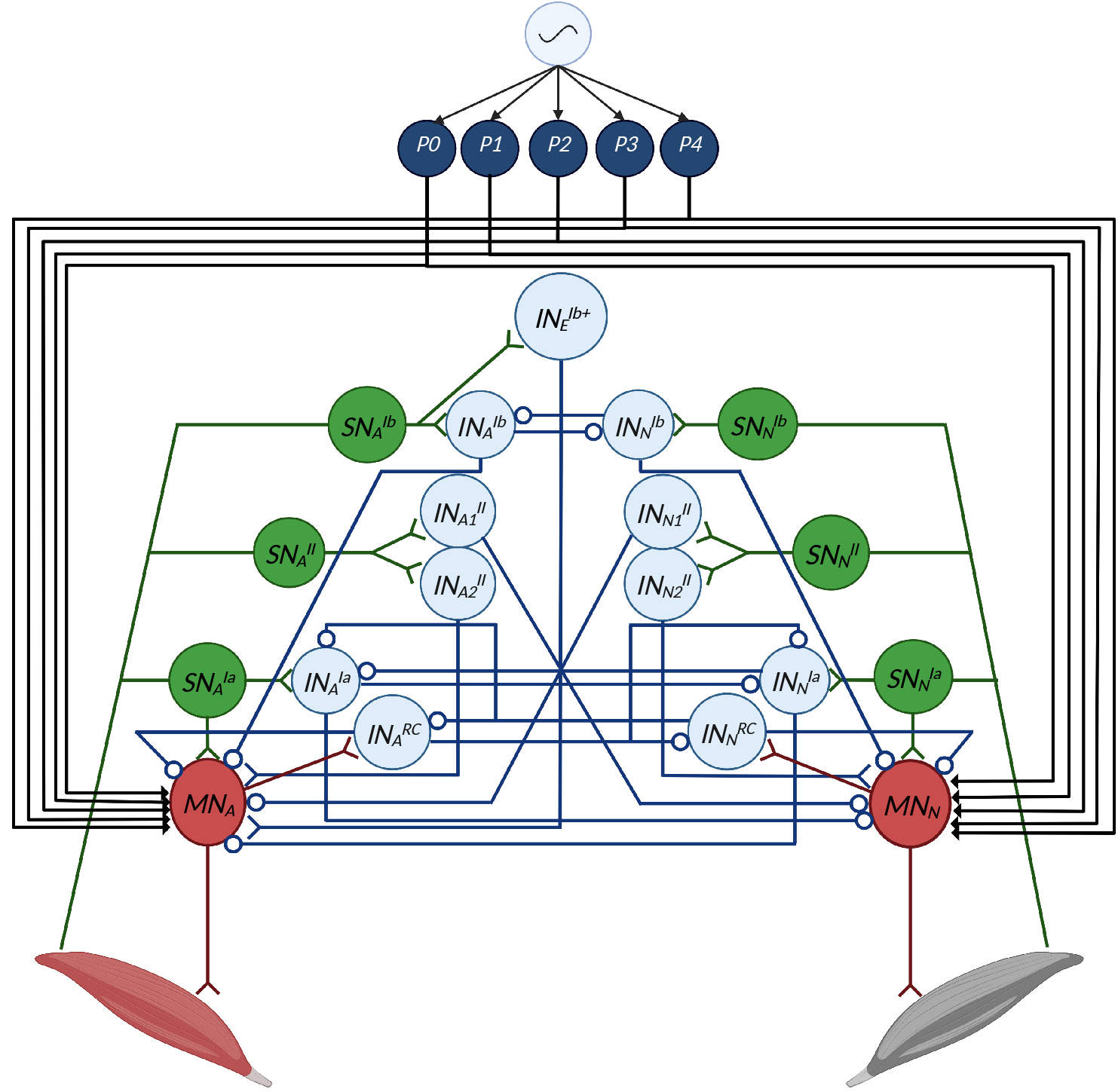
Spinal network between a muscle and its antagonist. The network includes reflexes driven by Ia, II, Ib afferents and Renshaw cells and inputs from CPGs’ patterns. Connections from patterns to motoneurons are represented by back arrows since these connections be both inhibitory and excitatory. ILPSO, GMAX, and HAMS also receive inputs from the balance controller. (Created with BioRender.com)

#### 2.3. Optimization process

In total, the controller’s parameters are 256, accounting also for 16 additional parameters regulating initial positions and velocities of the model’s DoFs. Because of the large size of the parameters’ space and the difficulties in obtaining a stable solution when the network is in an arbitrary state, the optimization process is divided into three steps: imitation objective, optimization for stability, and optimization of metabolic energy. In the first stage, we try determining the network’s parameters such that the output of neurons is within a plausible range and motoneurons’ activity resembles normal gait solutions. To achieve this, we begin with a previously obtained stable gait simulation generated by a simpler controller [36]. Given that we know the whole state trajectories of the musculoskeletal system, we can compute the sensory afferent inputs required by the bio-inspired controller. Therefore, we can optimize for network parameters efficiently without numerically integrating the equations of the musculoskeletal system. We call this step imitation learning because we try to imitate a simulated behavior without yet producing dynamically consistent stable gaits. The*P4*optimization objective is defined as follows:

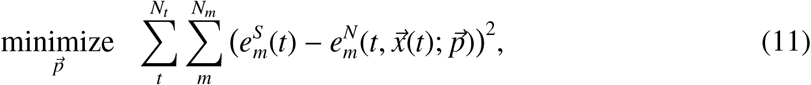

where 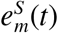 denotes the target simulated excitation of muscle *m* at time *t*, and 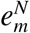he excitation of the network that depends on time *t*, the known state variables 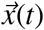, and parameters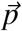. The above parameter solution does not produce stable gaits if we evaluate the model by numerically integrating the equations of motion. Our initial goal was to calibrate the network behavior within a reasonable range of operation in order to avoid neuron activities that are extreme and always make the model fall and optimization diverge. In fact, the imitation is only done to obtain a first usable solution for further optimization.

The second optimization aims at obtaining dynamically consistent stable gaits. To do so, we start integrating numerically the equation of motion producing dynamical gaits by minimizing the distance between the model’s and reference’s states as already expressed in equation 11, penalizing unstable falling solutions and solutions outside the desired range of speed, minimizing metabolic effort and joint limit torques. With this process, we aim to obtain stable solutions generated with our bio-inspired controller with physiological kinematics and muscle activation. The optimization is done using a CMA-ES algorithm with parameters λ = 40 and σ = 5 [23]. The cost function for this optimization is defined as follows:

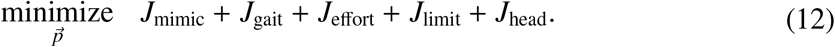

The term *J*_mimic_, represents the model mimicking the reference states as expressed in equation 11, *J*_gait_ penalizes the solution where the center of mass velocity is outside the [*v*_*min*_, *v*_*max*_] range (1.10-1.25 m/s for healthy human gait at normal speed) and the falling solutions. The model is considered to fall when the ratio between its center of mass height (*h*_*COM*_) to the initial state (*h*_*COM,i*_) is smaller than a termination height threshold set to 0.8 (^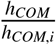^< 0.8). The term *J*_effort_ defines the rate of metabolic energy expenditure [47] normalized by the product of body mass and distance traveled. *J*_limit_ is associated with joint minimization of soft joint limit torques at the knee and ankle joints in order to avoid excessive joint angles [14]. Finally, *J*_head_ helps to maintain head stability by minimizing horizontal and vertical head accelerations outside the following ranges: [−4.90 − 4.90]*m*/*s*^2^ in the vertical direction, and [−2.45 − 2.45]*m*/*s*^2^ in the horizontal direction, as previously done by Ong et al. [36]. Concerning the weights, we assigned *w*_mimic_ = 10, *w*_gait_ = 100, *w*_effort_ = 1, *w*_limit_ = 0.1, and *w*_head_ = 0.25 in order to promote mainly stability and mimicking. Following this optimization, we use the resulting stable solution as initial condition to find the optimal that minimizes metabolic energy. To do so, we remove the mimicking component of the cost function and optimize for

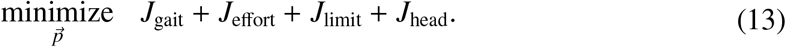

*J*_gait_ allows the stability of future explored solutions and *J*_effort_ allows convergence toward gait efficiency. In addition, we apply external perturbations to the pelvis and randomized internal perturbations to muscle excitation to obtain more robust and stable gaits. The external perturbation is a force of 100 N applied in the forward and backward direction for a duration of 0.2 s respectively after 3 s and 4 s after the beginning of the simulation. The internal perturbations are applied to sensory receptors. For each controller timestep, a random white Gaussian noise is sampled from a normal distribution with a standard deviation of *s* ∗ *noise*_*p*_, where *noise*_*p*_ is the proportional standard deviation of the normal distribution, and *s* is the perturbed sensory signal.

This three steps optimization process was used only to find a proper local optimum to replicate human gait behavior with a high number of parameters tuning the bio-inspired controller. However, once the local optimum is found, different gait behaviors can be reached starting from this solution by only optimizing according to equation 13 with the different gait behaviors targeted by *J*_gait_. These experiments are explored in the following section.

#### 2.4. Gait modulation

To study the capability of the proposed bio-inspired controller to reproduce different gait behaviors in human locomotion, we focus mainly on the modulation of locomotor speed. In this way, we aim to evaluate our controller’s performance, checking the maximum and minimum speeds it can achieve. Additionally, we evaluate gait analysis and muscle activation for three selected solutions far from the extremes of the achieved speed range since very slow or very fast speeds are more subject to producing artifacts in gait simulations. Therefore the three solutions selected are at 0.6 m/s, 1.2 m/s, and 1.6 m/s representing slow, intermediate, and fast speeds, respectively. To do so, we modulate the optimization parameters [*v*_*min*_, *v*_*max*_] in *J*_gait_. Furthermore, we use the data acquired from our model to have possible insights into the contribution of CPGs and spinal reflexes in the neuromotor control of human locomotion and gait modulation. To do so, first, we evaluate the inputs to motoneurons from CPGs and reflexes and how these affect the motoneurons’ output at different speeds. We then performed additional optimization where either CPGs parameters or reflexes parameters were fixed to investigate the modulation capabilities of each controller component. The fixed values of parameters are extracted from a reference solution of the model walking at 1.17 m/s with 0.79 m of step length and 0.67 s of step duration. This solution is the one where the optimizer converged without imposing any restriction on the target speed. Finally, we investigate which parameters majorly contribute to gait modulation for the three controller configurations: full control, fixed reflexes, and fixed CPGs. These parameters are identified through a correlation analysis with gait speed, step length, and step duration, were parameters that have a high level of positive or negative correlation with these three gait characteristics (above 0.80 in absolute value) are considered the potential major contributors to gait modulation [14]. The correlation analysis is conducted over 8 samples obtained through different target optimizations for each of the 3 controller configurations.

## 3. Results

Figure 7 shows the controller’s performance. When minimizing the cost function without imposing restrictions on gait speed, the model converges to a gait at 1.17 m/s of speed, 0.79 m step length, and 0.67 s step duration. Figure 6a shows qualitatively the different positions of the model’s joints through the gait cycle. The simulated pelvis tilt, hip flexion, knee angle, and ground reaction forces (GRFs) shown in Figure 6b faithfully represent the experimental observations from Schwartz et al. [42] illustrated by the shaded grey areas. Some discrepancies can be observed for the ankle angle that tends to have excessive dorsiflexion and lacks proper plantarflexion during push-off compared to experimental observations. Indeed, the ankle angle mostly maintains its values above the zero level of plantarflexion/dorsiflexion. This likely depends on the weak activation of gastrocnemius and soleus observed in Figure 6c. These muscles maintain a peak activation of 0.3 for the soleus and 0.2 for the gastrocnemius. However, the simulation replicates the temporal activations observed in experiments from Perry and Burnfield [37] for TA, GMAX, VAS, GAS, SOL, and HAMS. Concerning ILPSO, the muscle is active also outside its time range, having a consistent activation also in preswing. The model converges to different behaviors compared to experimental results for BFSH and RF that are active at the beginning and at the end of the gait cycle, respectively, rather than during swing.

**Figure 6:**
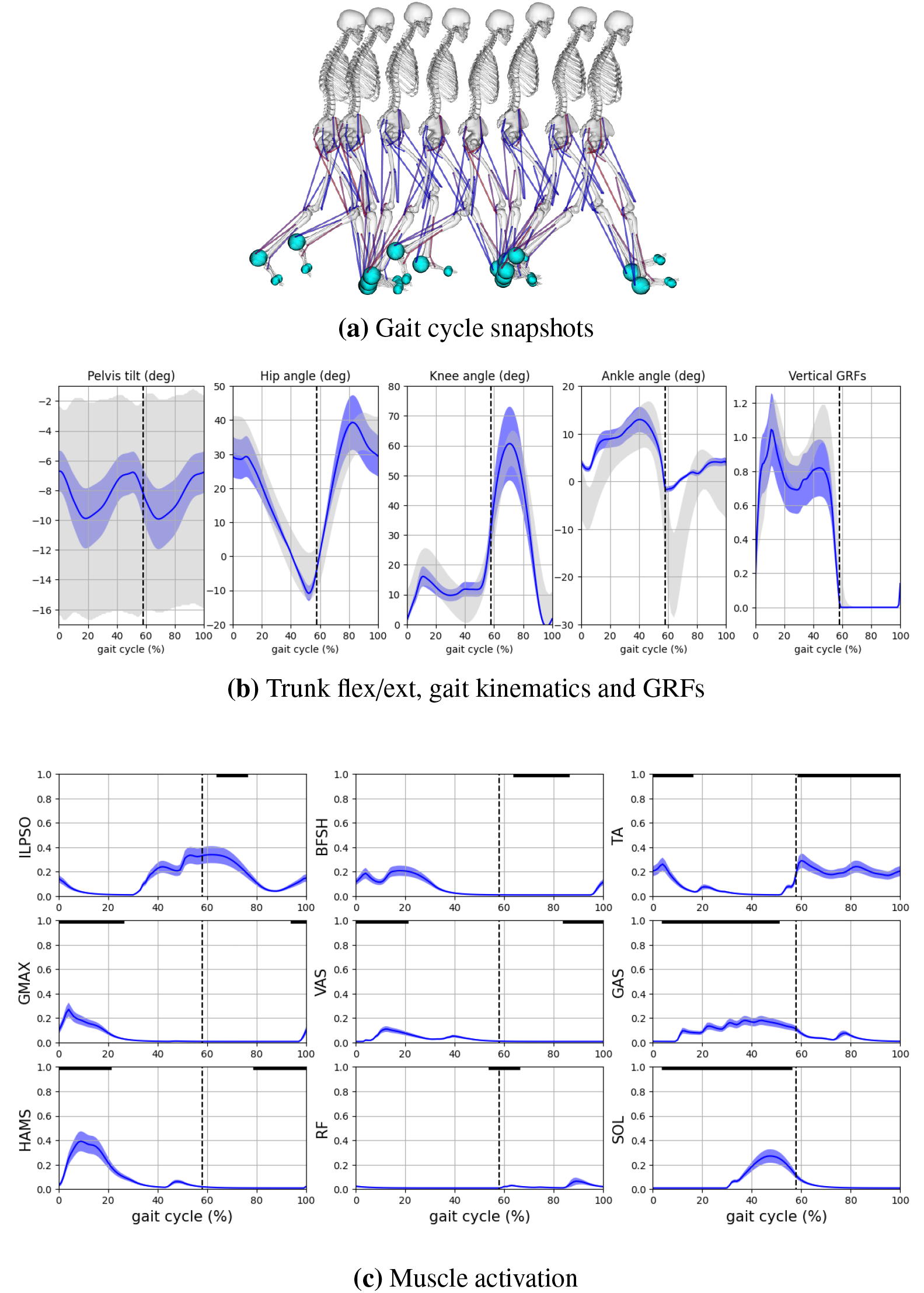
Gait analysis of simulated gait at 1.17 m/s. (6a) Model position at different times of the gait cycle. (6b) Kinematics and GRFs compared to experimental data [42]) grey areas report the observed experimental ranges for pelvis tilt, hip flexion, knee flexion, ankle dorsiflexion, and vertical GRFs. (6c) Muscle activation analysis: muscle activity over a gait cycle for the 9 muscles along the gait cycle. Blue curves represent the means of the gait signals through the gait cycles and the shaded areas the standard deviations. The activation curves are compared with the activation timing observed experimentally [37] and represented by the solid black lines on the top of the graphs.

**Figure 7:**
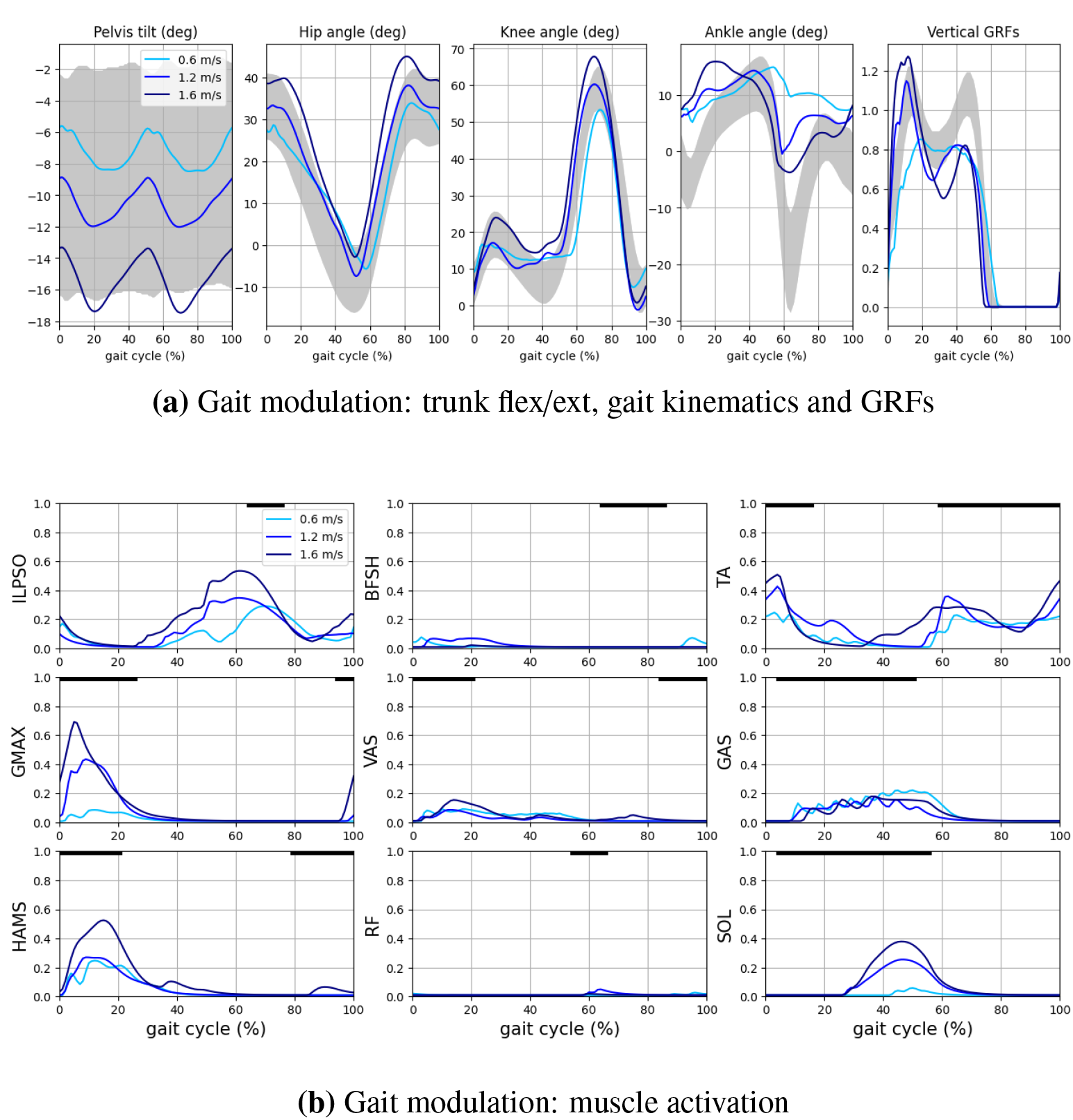
Gait modulation at 0.6, 1.2, and 1.6 m/s. (7a) Comparison of kinematics and GRFs among the three speeds. (7b) Changing muscle activation at different speeds. The activation curves are compared with the activation timing observed experimentally and represented by the solid black lines on the top of the graphs.

### 3.1. Gait modulation

By optimizing the controller’s parameters, the model could reproduce gaits from 0.3 to 1.9 m/s. Figure 7a shows the modulation of gait kinematics and GRFs at 0.6, 1.2, and 1.6 m/s representing slow, intermediate, and fast gaits, respectively. As speed increases, the pelvis tilt and the lean angle of the trunk increase in the forward direction by 8 degrees, and the hip flexion oscillates between 35 and -5 degrees at slow speed and 45 and -3 degrees at fast speed. Increasing amplitudes of knee flexion are also observed at high speed, having the peak flexion in swing of 53 degrees at 0.6 m/s and 68 degrees at 1.6 m/s. Fast speed also presents a consistent increase of ankle plantarflexion to -3 degrees of the ankle angle during ankle push-off, whereas this value is maintained at around 10 degrees of ankle dorsiflexion at slow speed. Concerning GRFs, the characteristic double peak shape is very weak at 0.6 m/s. Double peak amplitudes increase with the increase of speed, especially the first peak that shows the reaction with the impact with the ground during heel strike. The duration of the stance phase is reduced from 65% of the gait cycle at 0.6 m/s to 55% at 1.6 m/s. The behaviors of kinematics and GRFs modulation presented resemble the ones observed experimentally by Schwartz et al. [42]. Some differences are observed with the level of ankle dorsiflexion since the model tends to converge to a high-level of dorsiflexion that can differ from experimental data by 7 degrees during heel strike and by 12 degrees during push-off at slow and fast speeds. Additional differences are observed for the level of hip extension at slow speed and knee extension at high speed. In fact, Schwartz et al. observed that the maximum hip extension decreases at low speeds, and knee extension during stance increases at high speeds. In contrast, the model reproduced increased knee flexion during stance at high speeds and a similar level of maximum hip extension at 0.6 m/s compared to 1.2 and 1.6 m/s.

Muscle activity is affected by gait modulation mainly through the increase of activation with the increase in speed. In Figure 7b, ILPSO, GMAX, HAMS, TA, and SOL are the muscles that more consistently present an increment in muscle activation. TA and HAMS pass from a maximum activation of 0.2 at slow speed to 0.5 at fast speed, whereas ILPSO has a similar maximum activation at fast speed and a higher activation of 0.3 at slow speed. GMAX has the highest increment of muscle activation, passing from a maximum activity of 0.1 to 0.7. SOL also presents a consistent increase in its activity, passing from a value smaller than 0.1 at slow speed to 0.4 at fast speed. A lower increase is present for VAS at high speed, whereas no consistent variation in muscle activity can be observed for BFSH, RF, and GAS. Therefore, the increased plantarflexion with speed mainly depends on the increased activity of the soleus. In general, the activation amplitude of all muscles increases with speed, as observed experimentally by Cappellini et al. [8].

### 3.2. Gait modulation: CPGs and reflexes

Past controllers suggested spinal reflexes may be sufficient to generate human locomotion in simulations [18, 36]. These reflexes were regulated by state-machine mechanisms. We tested our controller to check whether the spinal connection implemented could generate rhythmic locomotion without the state-machine regulation or the presence of CPGs. With the removal of CPGs, the remaining parameters to optimize are 208. Even if the dimensional reduction could in principle simplify the convergence to a stable solution, no rhythmic gait could be reproduced, suggesting the need for the CPG networks to provide rhythm and timing in the absence of a state machine activating sensory feedback commands at specific times of the gait cycle. The simulation resulting from removing CPG parameters led to the human model in a standing position with the right leg in front of the left leg. The reflexes could generate the muscle activation necessary to maintain this position until the balance controller failed to stabilize the trunk, causing the model to fall. Dzeladini et al. [15] suggested that the tuning of CPGs applied only to hip muscles could easily modulate human locomotion where other muscles were controlled by sensory feedback. In order to test this hypothesis in our model, we set to 0 the CPG parameters for all muscles except hip muscles and re-optimized the parameters according to the step explained in section 2.3. The resulting simulation showed very similar behavior to the one obtained without any CPG parameters, suggesting CPGs inputs may have an important contribution also for knee and ankle muscles. It should be noted that our CPG model provides only a rough waveform (made of the 5 primitives), while Dzeladini’s CPG provides a detailed waveform replicating the sensory-driven control signals.

To investigate the contribution of each controller component in gait modulation, we investigated the inputs from CPG circuits, spinal reflexes, and balance controller provided to the motoneurons. Figure 8 shows how these signals contribute to generating motoneurons inputs and outputs following equation 3. Generally, in the model, the net effect of the reflex circuits tends mainly to inhibit the motoneurons providing a negative stimulation through the whole gait cycle with the exception of ILPSO and TA. Reflexes also facilitate the activation of VAS and GAS during swing for all the speed ranges and the activation of BFSH and HAMS at slow speeds. Instead, for each muscle, CPGs present specific regions of the gait cycle where they excite or inhibit the motoneuron. In this regard, CPGs prevent the activation of specific muscles in specific cycle phases, such as VAS and GAS in swing that were stimulated by the reflex circuits. CPGs’ patterns tend to increase the amplitude of inhibition or excitation with increasing speed. This is especially the case for SOL, where the growing muscle activation with speed is primarily due to the increased excitation from CPGs circuits. CPGs activity also helps to have a consistent muscle activation of ILPSO in swing, but it tends to increase the activity at slow speed, and the lower muscle activity in swing is achieved by reflexes that inhibit ILPSO during swing at 0.6 m/s. The balance controller is applied only to ILPSO, GMAX, and HAMS, and seems to be the leading cause of HAMS activation since the CPGs excitation is entirely inhibited by spinal reflexes.

**Figure 8:**
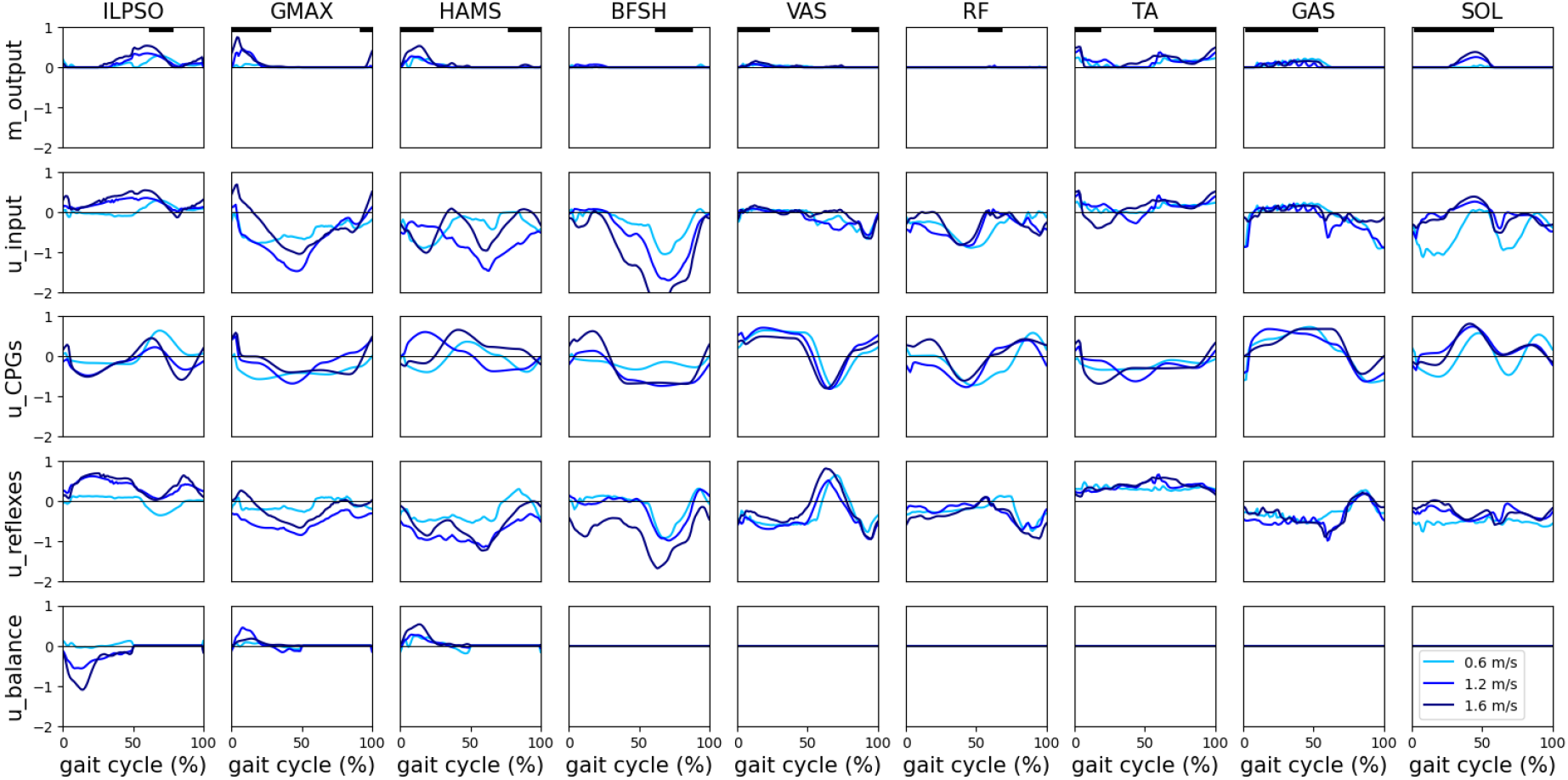
CPGs, reflexes, and balance inputs at 0.6, 1.2, and 1.6 m/s. CPGs, reflexes, and balance generate motoneurons inputs and outputs according to equation 3. The contribution of the three controller components is compared through the three speeds selected.

Additionally, we performed optimizations for different target speeds by keeping fixed reflexes parameters or CPGs parameters. Table 2 compares the achieved ranges of speed, step length, and step duration for the controller optimizing all parameters (full control), maintaining reflexes parameters fixed (fixed reflexes), and maintaining CPGs parameters fixed (fixed CPGs). The optimization of all parameters allows reaching wide ranges of speed from 0.30 to 1.86 m/s with small and large step lengths (0.23 to 1.08 m) and step durations (0.53 to 0.84 s). Removing reflexes parameters’ optimization allows reaching ranges similar to the ones obtained in full control. However, the missing optimization of CPGs parameters significantly limits the controller’s capabilities to modulate step duration, passing from a range covering 0.53-0.84 s to 0.64-0.67 s. Consequently, the optimization tends to achieve slow or fast speeds, mainly modulating the step length to reach large values of 1.21 m at high speed. The achieved value of step length is higher than the one in full control because the model converges to a more energetically efficient gait reducing the step duration when all parameters are optimized. By maintaining only the oscillatory frequency fixed and optimizing all the other parameters, the controller covered a similar range compared to the configuration with fixed CPGs, suggesting that the CPG frequency could be the principal modulator of step duration. However, we verified that the modulation of CPG frequency alone is insufficient to converge to different gait behaviors. Indeed, the model loses stability without significant changes in gait speed when only the CPG frequency is tuned. This result implies that the modulation of CPG frequency alone may be necessary but not sufficient to modulate step duration.

**Table 2:** Evaluation of speed, step length, and step duration ranges achieved by the bioinspired controller in 3 configurations: full control where all parameters are optimized, fixed reflexes where all reflexes parameters are fixed, and fixed CPGs where all CPGs parameters are fixed.

|  | Full control | Fixed reflexes | Fixed CPGs |
| --- | --- | --- | --- |
| Speed (m/s) | [0.30-1.91] | [0.30-1.91] | [0.44-1.91] |
| Step length (m) | [0.23-1.08] | [0.23-1.08] | [0.30-1.21] |
| Step duration (s) | [0.53-0.84] | [0.56-0.84] | [0.64-0.67] |

#### 3.2.1. Correlation analysis

The correlation analysis reported in Table 3 gives indications on which parameters had a correlation higher than 0.8 with the main gait characteristics and, therefore, those that could be the main responsible for gait modulation in the three controllers configurations. Specific parameters are identified as:

**Table 3:**
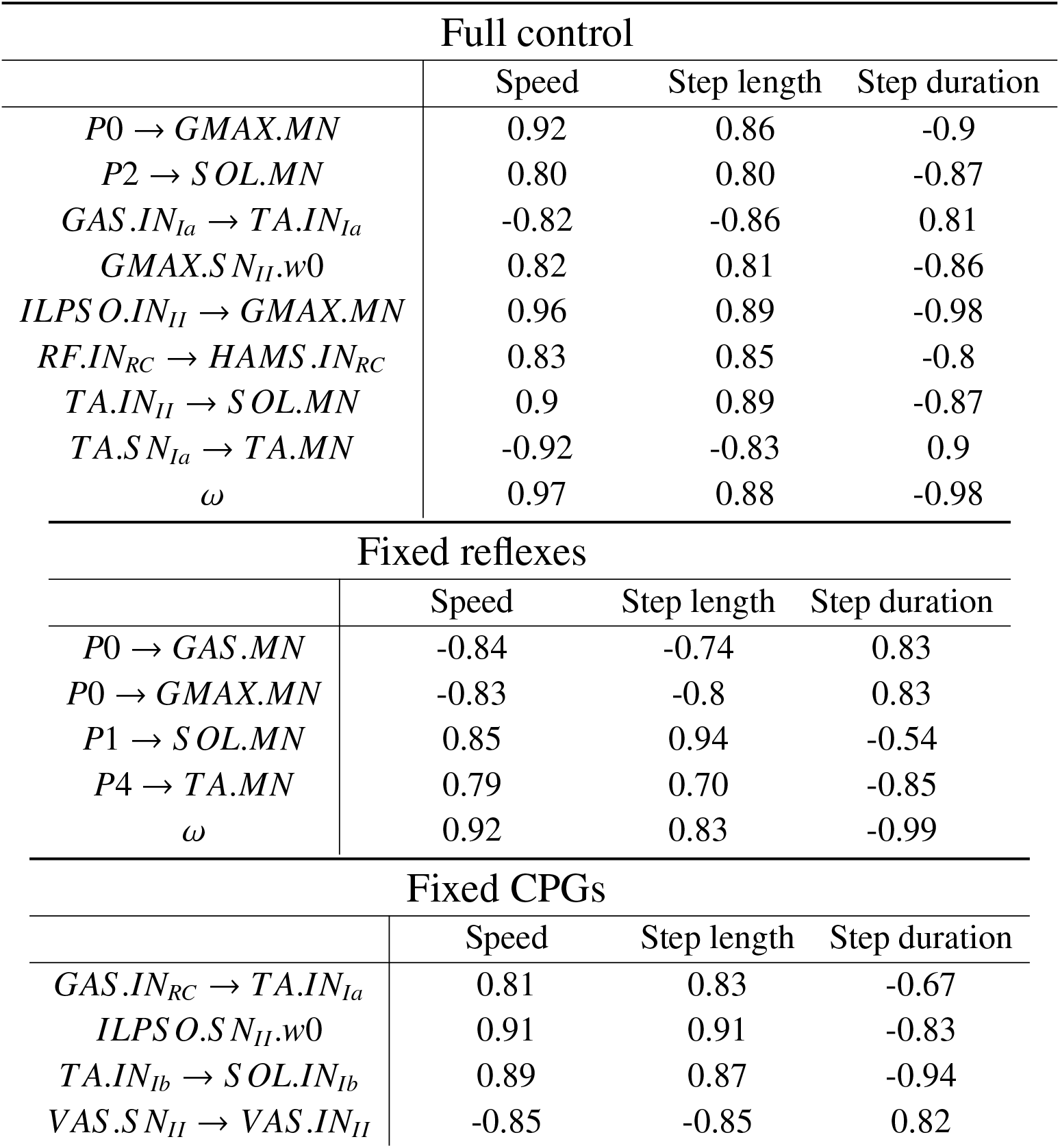
Correlations coefficients of controller’s parameters contributing to the modulation of speed, step length, and step duration in the 3 controller’s configurations: full control, fixed reflexes, and fixed CPGs.

- *Pk* → *M*.*MN* for the input pattern *Pk* weighted connections to motoneuron *M*.*MN*, where *M* is the muscle name.
- ω for phase oscillator frequency.
- 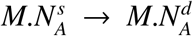 for parameters regulating the weighted synaptic connections between the source neuron of a specific muscle 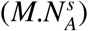and the destination neuron of the target muscle 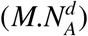. *N* represents the type of neuron and can be either *S N, IN*, or *MN*, and *A* represents the type of afferent and can be either *Ia, II, Ib, Ib*+, or *RC*
- *M*.*N*_*A*_.*w*0 is the activation offset of the neuron *N*_*A*_ regulating the neuronal response.

In full control, both reflexes and patterns’ connections seem to contribute to gait modulation. The first (P0) and third (P2) patterns connections to extensor muscles like GMAX and SOL positively correlate with increasing speed and step length and decreasing step duration.

The CPGs’ frequency (ω) has a highly consistent correlation with gait speed and step duration, suggesting again the direct influence of this parameter on gait frequency. The only reflex parameter representing an excitatory connection is the monosynaptic excitation of Ia afferents from TA (*TA*.*S N*_*Ia*_ → *TA*.*MN*), having a negative correlation with speed and favoring increased dorsiflexion during slow gaits. Another relevant parameter is the length offset of the II somatosensory neuron of GMAX (*GMAX*.*S N*_*II*_.*w*0) with a positive correlation with speed and step length meaning a higher level of stretch is needed to activate length feedback from II afferents. The other reflex parameters presented are inhibitory connections, which implies that a highly negative correlation with a gait characteristic (either speed, step length, or step duration) means an increased inhibition with the increase of that gait characteristic. The II interneuron of ILPSO tends to decrease its inhibition to GMAX motoneuron (*ILPS O*.*IN*_*II*_ → *GMAX*.*MN*) when speed increases and step duration increases, favoring the activation of GMAX in these conditions. The same mechanism is involved in facilitating the activation of SOL through the decreasing inhibition from II interneuron of TA (*TA*.*IN*_*II*_ → *S OL*.*MN*). The last two reflex parameters for the controller in full control configuration involve the reciprocal inhibition mechanisms of interneurons and Renshaw cells.GAS’s Ia interneuron increases the inhibition to TA’s Ia interneuron with increasing speed (*GAS* .*IN*_*Ia*_ → *TA*.*IN*_*Ia*_), enhancing the activation of GAS itself because of the decreased inhibition from *TA*.*IN*_*Ia*_. Then, the Renshaw cell of RF decreases its inhibition to the Renshaw cell of HAMS (*RF*.*IN*_*RC*_ → *HAMS* .*IN*_*RC*_) with increasing speed, favoring the inhibition of the hamstrings muscle. Indeed, from the previous analysis, the increased activation of HAMS with speed was mainly due to the input from the balance controller.

Concerning the configuration with fixed reflexes, CPGs’ frequency (ω) highly correlates with speed and step duration also in this case. Another modulator for step duration is the input from the fifth pattern to TA’s motoneuron (*P*4 → *TA*.*MN*), which indeed increases its activation at the end of the gait cycle with increasing speed. The first pattern (*P*0) tends to increase the inhibition to GAS and GMAX at the very beginning of the gait cycle with increasing speed. Then, speed modulation through modulation of step length is enhanced by tuning the excitation from the second pattern (*P*1) to SOL motoneuron (*S OL*.*MN*) in order to increase propulsion in stance.

When CPGs parameters are fixed, speed modulation happens mainly through step length changing because of the controller’s limited capability to modulate step duration without tuning CPGs’ frequency. The controller tends to increase step length by increasing the offset to enhance the length feedback of ILPSO (*ILPS O*.*S N*_*II*_.*w*0). II afferents are also involved with the decreased length feedback of VAS muscle with speed rising through the excitation of *VAS* .*IN*_*II*_ from *VAS* .*S N*_*II*_. The last two relevant parameters concern the inhibitory connections of Renshaw cells and Ib afferents. The Renshaw cell interneuron of GAS (*GAS* .*IN*_*RC*_) decreases its inhibition to the Ia interneuron of TA (*TA*.*IN*_*Ia*_), decreasing the activation of GAS itself at fastest speeds. Higher speeds should, in principle, increase the activation of GAS, but in the modulation of muscle activation, we previously observed that the optimizer tends to maintain the same activation level for the gastrocnemius muscle during speed modulation. Finally, the Ib inhibitory interneuron of TA (*TA*.*IN*_*Ib*_) decreases its inhibition to the Ib interneuron of SOL, allowing the inhibition of this muscle. Indeed, we previously observed that the increased muscle activation of soleus at higher speeds was not due to the input from spinal reflexes but primarily due to increasing excitatory inputs from CPGs.

## 4. Discussion

In this study, we aim to investigate the possibility of controlling human locomotion by relying only on spianl reflexes not regulated by a state machine mechanism and to investigate the contribution of both CPGs and spinal reflexes in generating locomotor output. To do so, we developed a bio-inspired controller composed of a balance controller, a CPG network, and a sensory feedback network based on physiological spinal reflexes maintaining a state-machine mechanism only for the balance of the trunk. The proposed controller could replicate human kinematics and ground reaction forces (GRFs) with some limitations in the ankle angle in which the model converges to an excessive dorsiflexion behavior. Regarding muscle activation, the model could reproduce most of muscle activation timings observed experimentally, with the exception of BFSH, which is active outside its range in human recording. The proposed network could probably generate muscle activation closer to physiological activity with additional optimizations. However, finding this global optimal solution results challenging because of the large number of parameters.

Many aspects of speed modulation from human recordings, such as the increased amplitudes of flexion/extension movements and the increased muscle activation with growing speed, are also matched by the model. Concerning the role of CPGs and spinal reflexes in the neural control of human movement, we investigated the possibility of finding stable solutions without relying on CPGs as suggested by previous neuromechanical studies [18, 36]. In our optimizations, we could not find any stable rhythmic behavior in the absence of CPGs’ commands even if the number of parameters to optimize significantly decreased, suggesting the need for the CPG network to provide rhythm and timing in the absence of a state machine activating sensory feedback commands at specific times of the gait cycle. Therefore, reflex-based circuits are always active through self-regulation by the afferents, lacking any timing information without CPGs. Similarly, a pure CPG network without reflexes leads to unstable solutions. While we cannot rule out that a different network topology might give rise to high-quality gaits, our model highlights the need for both types to achieve stable and natural movements. Indeed, natural locomotor behavior emerges when both CPGs and spinal reflexes are active. Our study suggests that the state-machines used in previous sensory-driven models [19, 36] could in fact be replaced by CPGs and that one of the main roles of CPGs, in addition to simplifying speed control [15], is to serve as gating mechanism that ensures that reflexes do not affect muscles all the time but only at specific moments of the locomotor cycle.

More specifically, the performances on gait modulation while either reflex circuits or CPGs commands were fixed, and the corresponding correlation analysis highlighted the importance of CPGs’ frequency in changing the step duration. Therefore, in the model, CPGs have a crucial role in determining gait timing. Additionally, the analysis of neural inputs to motoneurons showed that the net inputs of reflexes are mainly inhibitory through the gait cycle for the proposed model, except for ILPSO and TA, which globally receive excitatory inputs. CPGs’ patterns excite or inhibit motoneurons in specific phases of the gait cycle to allow or prevent muscle activation. Therefore, CPGs seem to be important to determine activation timing other than gait frequency. Such a control strategy is similar to the one proposed by Laquaniti et al. [28] where the timing and magnitude of EMG activity are tuned via proprioceptive feedback and CPGs that control the basic rhythms and patterns of motoneuron activation. However, it should be highlighted that the five locomotor primitives described by Ivanenko et al. and Laquaniti et al. [25, 28] were not equally spaced in the gait cycle phase as they are in our controller. This is because, in these studies, the primitives were extracted with factorization of EMG activity. Yet, this activity is the result of the global input received by muscles without being able to distinguish which input was coming from spinal reflexes and which one from CPGs circuits. Therefore, we decided to simplify the distribution of the five primitives and equally space the patterns through the gait cycle since the primitives measured in experiments could hardly be generated by the CPGs commands alone. This choice still leads to largely reproducing the experimental activation timing.

The modulation of gait reflexes alone could still regulate muscle activation to achieve different gait behaviors, mainly through the modulation of step length. The correlation analysis highlighted the possible parameters responsible for this behavior, such as the offset of II fibers regulating the level of stretch necessary to activate length feedback for ILPSO and GMAX. Indeed, increasing these parameters allowed larger amplitude for hip flexion/extension, promoting larger step lengths.

In general, the proposed controller presents a highly redundant system where several different combinations of neural inputs can generate the same muscle activation. The correlation analysis gave possible insights into which parameters could be the most relevant in the control of gait modulation. Yet, given the high redundancy, a separate and more extensive study would be necessary. Possibly, this study should include a large dataset of optimizations and additional elements of the cost function that could guide toward the best combination of neural inputs to generate specific muscle excitation, such as the minimization of the total neuronal activation. Then, the results should ideally be validated by experimental measurements.

Some limitations of the proposed controller should be considered. Because of the large number of parameters, finding a stable solution replicating human walking with the proposed controller may be challenging since it requires the three optimization stages described in section 2.3. However, once this solution is found, it can be used as a starting point to explore different gait behaviors by only performing the last optimization stage. In this way, we could reproduce a wide range of speeds comparable to or larger than the ones previously obtained by other neuromechanical controllers [43, 36, 14]. Yet, it should be highlighted that the initial stage of the optimization requires the imitation objective from a previous solution found with a different neuromechanical controller. This step was necessary because some parameter combinations can quickly saturate the neurons’ output and overexcite or excessively inhibit the network resulting in permanent or no muscle activation and leading to model failures during volatile movements. There is a low probability that a random initialization of parameters can make the convergence to a stable solution. If the model falls initially, it is hard to learn a good solution to improve the gait and escape the local minimum. However, the use of the imitation objective implies that any lack of performance from the imitated solution in replicating human movement will probably reflect a lack of performance of the bio-inspired controller. This has probably been the case for the excessive dorsiflexion behavior performed by our model since many solutions of the reflex-based controller proposed by Ong et al. [36] that we used as imitation objective presented the excessive dorsiflexion behavior. Therefore, the proper choice of the initial imitation objective is crucial for the correct optimization of our model.

Further considerations should also be made for the design of the reflex controller. In paragraph 2.2.3, we explained how we simplified the expressions for the sensory receptors to capture the general trend and prevent an excessive number of physiological parameters. In reality, the dynamics of these receptors are very complex [32, 33], and there is little evidence why the same model identified in specific animal experiments can generalize to humans in the presence of dynamic movements.

Despite these limitations, the bio-inspired controller we propose is a promising tool for investigating spinal circuits in human locomotion. Indeed, we have already shown the insights this model could give into the relationship between CPGs and spinal reflexes. Further suggestions could be provided in investigating pathological gaits. Past studies tried reproducing neural pathologies with neuromechanical simulation by extending previous controllers, including specific connections to model the pathology in the desired degree of freedom [6]. However, the controller proposed could be more suitable for studying neuropathologies like hyperreflexia considering the effects of both excessive inputs from Ia fibers and the lack of reciprocal inhibition. Furthermore, further aspects of gait modulation regulating standing to walking transitions and acceleration and deceleration mechanisms can be investigated. Additionally, this controller could be used as a starting point to further extend the modeling of the neuromotor system by including the implementation of additional spinal neural connections like γ-motoneurons [16] and descending inputs from the brainstem and other supra spinal brain areas, even though this would increase even more the controller’s complexity and the total number of parameters. Additionally, future implementations could include less abstract and more realistic CPG models, for instance, based on more detailed models previously proposed for mammalian circuits that could potentially be taken as a reference for modeling human locomotion [5, 4, 12, 11]. Additional connections between CPGs and spinal reflexes may be implemented, allowing somatosensory neurons to interact and modulate CPGs’ patterns and CPGs’ patterns to interact with spinal interneurons other than motoneurons.

## Conclusions

This study proposes a novel physiologically plausible neuromechanical controller maintaining a good balance between complexity and realism to investigate the spinal components governing human locomotion. The controller is composed of a balance controller from Ong et al. [36], a CPG network inspired by Aoi et al. [3], and a sensory feedback network that takes into account the main reflex connections in the spinal cord without being tuned by a state machine. The controller demonstrated the ability to reproduce key behaviors of human locomotion and its modulation in simulations. Results from optimizations suggested that rhythmic locomotion could not be achieved with the only contribution of spinal reflexes without accounting for a state machine mechanism. This suggests the possible need for CPG networks to generate rhythmic movements by guiding muscle activation timing in specific phases of the gait cycle. The modulation of either CPGs or reflexes parameters or both could reproduce wide ranges of gait behaviors, highlighting the high level of redundancy in human locomotor control. The modulation of CPGs’ frequency appeared to be crucial for regulating gait cycle duration. The proposed controller demonstrated to be a promising tool to provide many other indications on how the spinal cord may produce locomotor outputs.

## Supplementary materials

All the codes necessary to replicate our experiments and the parameters and files of our simulation can be found in https://github.com/DiRussoAndrea/Spinal_controller. The SCONE version containing the implementation of the proposed controller can be found in https://gitlab.com/simgait/SCONE.

